# HiddenVis: a Hidden State Visualization Toolkit to Visualize and Interpret Deep Learning Models for Time Series Data

**DOI:** 10.1101/2020.12.11.422030

**Authors:** Jingquan Yan, Ruichen Rong, Guanghua Xiao, Xiaowei Zhan

## Abstract

Accelerometers provide continuous measurements of human movements. Utilizing their data exhibits great potential for monitoring or predicting human health outcomes. In this direction, researchers are active in developing machine-learning algorithms. However, compared to understanding deep learning models, visualizing and interpreting such models is equally important but has been less well-developed. To address these limitations, we constructed an online software tool to visualize the hidden states inferred from the deep learning models and to compare different cohorts based on outcome-associated temporal profiles. We demonstrated the utility of the model using the National Health and Nutrition Examination Survey (NHANES) dataset. Using this tool, we discovered that two time periods of daily activity contribute to significant differences in 5-year mortalities. To disseminate the software to the broader community for the analysis of accelerometer data, we provide the work as open-source code at https://github.com/jyan97/HiddenVis.

## 1. Introduction

Time-series data is defined as data captured over the course of time and which often contain temporal dependencies. A powerful analysis framework for time series data requires the modeling of correlations between the measurements in closed time frames. Several neural network modules are suitable: Recurrent neural network (RNN), gated recurrent unit (GRU), and long short-term memory (LSTM)[1][2]. Among those, the LSTM model can naturally incorporate the information carriers in both the short time window and long time window. The temporal patterns are learned via recurrent connections in each RNN layer, and then such information is passed to higher layers. The temporal dependencies are modeled as hidden states, which are the representations of previous inputs. Due to the good performance of RNN models for time-series data, they have been gradually adapted in biomedical research to predict clinical events[3]. However, the black box nature of neural networks makes it hard to understand hidden states, and users are also unlikely to further explore the time series data with the temporal information that RNN studied at each step.

In this study, we aim to develop a tool for facilitating the understanding of the LSTM model, as the interpretation and understanding of potential features contributing to the final prediction performance are important in real-world application, in particular for biomedical and clinical research. Based on the LSTM models described above, we seek to identify the elements that contribute to successful predictions. This works poses several challenges. First, interpretive sequential models are less well studied than convolution models, as convolution models are often linked with pictures and easily interpretable by the human eye. Second, while some sequential models with transformer structures are studied[4], they often require large sample sizes that are not available in most medical researches. The closest work we found is VisLSTM, a visualization framework for unsupervised LSTM models. We extend its idea to study the hidden states in a supervised learning fashion. Our goal is to develop a web-based toolkit, Hidden State Visualization Toolkit (HiddenVis) to visualize and facilitate the interpretations of sequential models for accelerometer data. Additionally, HiddenVis can visualize the hidden states, match input samples with similar patterns and explore the potential relation among covariates.

An accelerometer is a wearable device that continuously collects movement statistics from the wearer. It can be used to monitor host health in both the short and long term[5]. As such, there is increasing interest in investigating accelerometer data and their associations to health outcomes[6][7]. Accelerometer data are unique as they have inherently strong autocorrelations due to the data collection procedure. Traditionally, time-series statistics have been used to analyze accelerometer data[8]. Recently, the advancements of deep learning (DL) models have shown promising performances[9]. However, the DL models usually have thousands to millions of model parameters, spanning multiple neural network layers. Understanding the mechanism of the model and interpreting the model outcomes have been challenging[10]. We thus are motivated to develop visualization software to overcome these limitations in the analysis of accelerometer data. We demonstrate the functions and usages of the HiddenVis toolkit by one specific example of modeling the data from The National Health and Nutrition Examination Survey (NHANES). It visualizes the hidden states of a deep LSTM-FC autoencoder network, reveals time frames associated with the 5-year mortality outcomes, and facilitates the stratification of the recruited cohort. We deliver the software as open-sourced code in GitHub (https://github.com/jyan97/HiddenVis). It is also widely applicable to other temporal models where hidden states can be interrogated in search of their potential associations with health outcomes.

## 2. Case Study to Demonstrate Functions of the Tool

### 2.1. Study and Data Description

The Centers for Disease Control (CDC) conducts the National Health and Nutrition Examination Survey (NHANES), a large, stratified, multistage survey study that collects health and nutrition data on the US population to assess the health and nutritional status of adults and children in the United States. The NHANES gathers interview data, including demographic, socioeconomic, dietary, health-related questions, physical examinations, medical, dental and physiological measurements, and laboratory tests for thousands of participants every year. Since 2003/2004, minute-level physical activity (PA) has also been assessed in NHANES using an accelerometer for each participant for seven consecutive days[6]. This device measures vertical acceleration, and accelerations are transformed into activity count values. In addition to interviews and PA data, NHANES has also collected survival outcome data for all participants.

In this study, we used data from participants enrolled in 2003-2004, 2005-2006. The exclusion criteria for data analysis are: (1) Younger than 50 or older than 85 at the time when they wore the accelerometers; (2) With fewer than 3 days with at least 10 hours of wear time for PA activity monitoring; (3) Survival outcome data is missing; (4) Follow-up time is less than 5 years. Based on these selection and exclusion criteria, 2,977 participants were included in the modeling and analysis. Though we have already excluded all the participants who have fewer than 3 days of data with at least 10 hours of estimated wear time, the remaining physical activity data still exhibits various missing patterns over days, so imputation is necessary for further analysis. There are 129 participants missing 4 days activity counts, 192 participants with 3 days missing, 316 participants with 2 days missing and 699 participants with 1 day missing. The remaining 1,642 participants have fully recorded physical activity counts.

In order to impute the missing data, we assume that physical activity patterns are similar during the weekend as during the week, and that consecutive days are more likely to have the same physical activity pattern than nonconsecutive ones. We implemented the following missing data imputation method: First, for each participant, if activity counts on both Saturday and Sunday are missing, we impute these two days with the nearest workdays. Second, if either of these two weekend days is missing, we impute the missing day with the other one. Finally, we impute each remaining missing day with its nearest predecessor day. After doing so, each participant has a total of 1,440*7=10,080 PA counts in 7 days.

### 2.2 Model description

Our goal is to use time-series physical activity data to predict 5-year survival outcomes (death or alive as a binary variable) of participants in the NHANES study. We employed a 4-hour sliding window with a 3-hour 20-minute stride to calculate the mean, max and standard deviation from the original 10,080 minutes activity count, and thus this gives us 51*3 summarized PA data for each participant. Now the original activity data is transferred to the size of 2,977*51*3 and ready to feed into our training model.

RNN is a simple and efficient choice to analyses the time series PA data. However, it has a deficiency where the gradient of some of its weights starts to increase or decrease drastically while training for too many steps. This is known as the vanishing gradient problem. Thus we use LSTM, an improved RNN model, instead of the original RNN model for analysis to fit the PA data to predict mortality.

Our training model is shown in Figure 1. It is an auto-encoder model where one LSTM with 3 recurrent layers serves as the encoder, one fully connected network for reconstruction (decoder) and one fully connected network for mortality prediction. The prediction and reconstruction loss are combined for further back propagation. The LSTM encoder has 40 hidden layers, which give us 40 features at each input in the hidden state. Therefore, for each participant, we have 40 hidden layer features for each time input, and 50 hidden states (*h*_0_, *h*_1_, …, *h*_49_) for one participant. For this LSTM model, we have a total of 2,977*50*40 hidden values for all 2,977 participants. Next, we will demonstrate how to visualize and explore these 2,977*50*40 hidden states with our toolkit.

**Figure 1.**
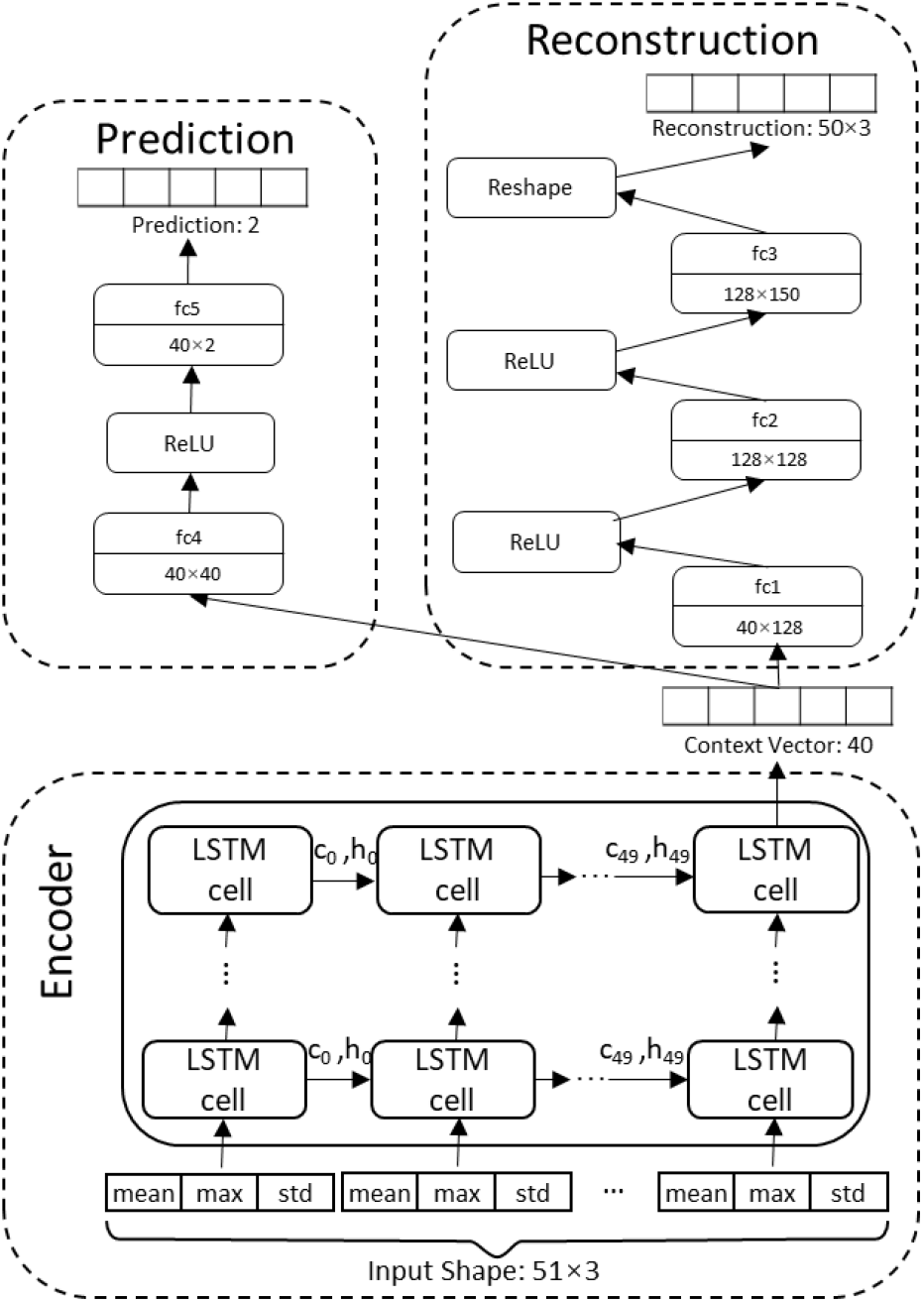
LSTM auto-encoder model

## 3. HiddenVis Tool Function Demonstration

### 3.1 Hidden State Visualization

First, we will show the user interface and a visualization plot for hidden states for a selected participant (or sample) in Figure 2. The SEQN parameter is the index number for each participant/sample. In Figure 2.a, the sample number (SEQN) indicates which sample has been selected to display the results in the chart. Figure 2.b displays the values for all hidden states (n=50) of all hidden layers (n=40) for the selected sample, and it shows how the values of each hidden layer vary over time. The X-axis represents the time-window and Y-axis represents feature values for each hidden layer over all 50 time-windows. Each curve stands for an individual hidden layer and is distinguished with different colors.

**Figure 2.**
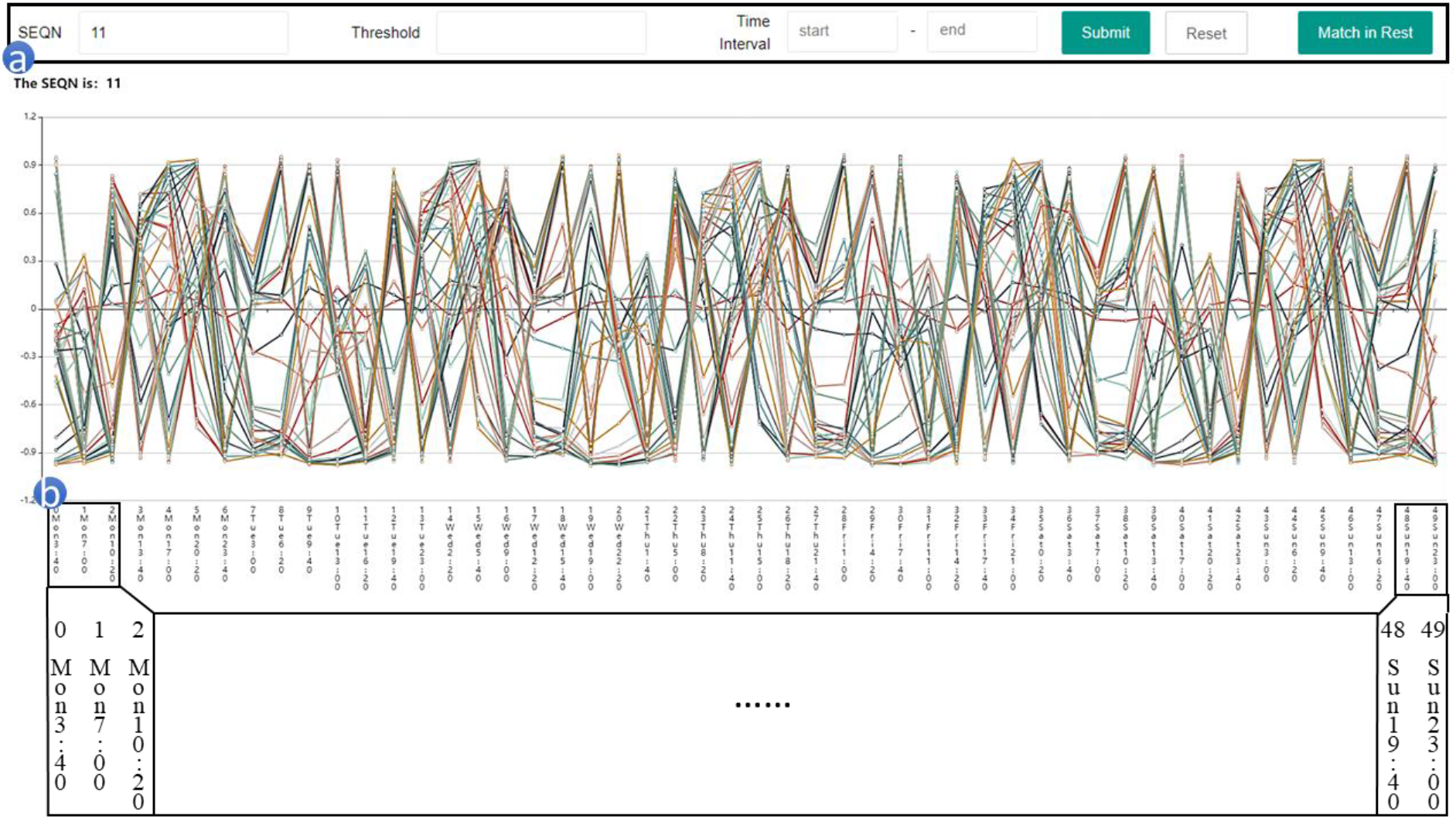
HiddenVis user interface and visualization

In addition to displaying all hidden layers for one participant using different colors, users can also specify specific selection criteria to select and highlight active hidden layers of interest. For example, Figure 3 displays all the hidden layers for participant #11 (same as in Figure 2), while only the hidden layers with values greater than threshold 0.2 during time interval 12 to 13 are highlighted in blue, and other layers that do not meet up the threshold requirement are marked in grey. The selection criteria are the combination of the threshold for feature values and with the time intervals, and those hidden layers that are larger than the threshold during the given time intervals are considered active hidden layers.

**Figure 3.**
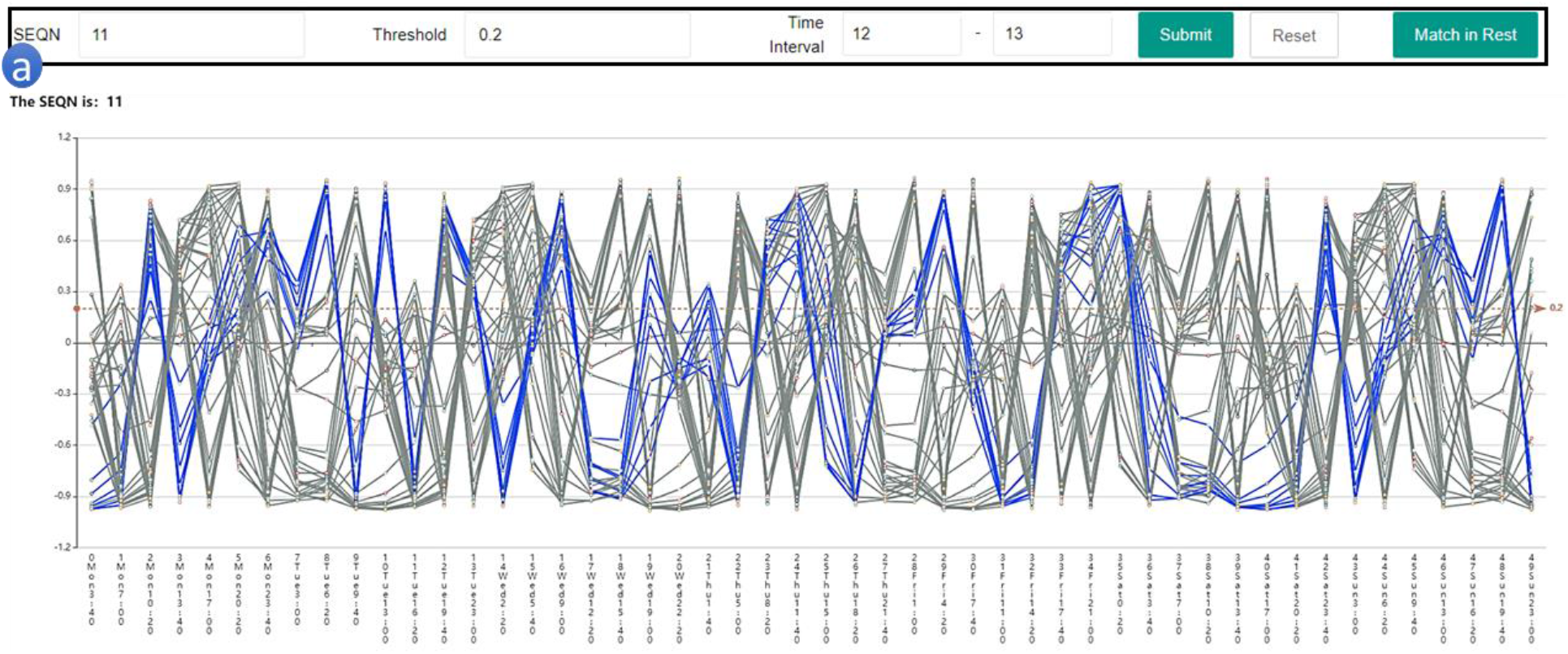
Hidden state visualization with threshold and time interval input

### 3.2 Identify Matched Samples

After visualizing hidden states, users can identify all samples with same hidden states patterns by clicking the “Match in Rest” button in Figure 2. We define the “same pattern” as those samples who share the same set of hidden layers that satisfy the given criteria (threshold and time interval). Formally, for one specific sample **s**_**1**_ with a hidden layer set ***H***, given an input criteria set 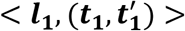 where ***l***_**1**_ stands for the threshold and the time interval is from ***t***_**1**_ to 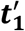, this criteria will filter a hidden layer set ***H′***, which is a subset of ***H***. Then we will match the remaining samples who meet this criteria set 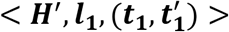. That is, for any layer in ***H′***, the layer must be larger than ***l***_**1**_ during time interval ***t***_**1**_ to 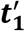.

The toolkit will return a chart (Figure 4) which contains all the matched samples and their corresponding clinical data. For comparison, we can check the box (Figure 4.a) in front of any sample of interest and click “Show the Chart” (Figure 4.b) to visualize the hidden layer flow chart and further compare the flow chart to the chart shown in Figure 3.

**Figure 4.**
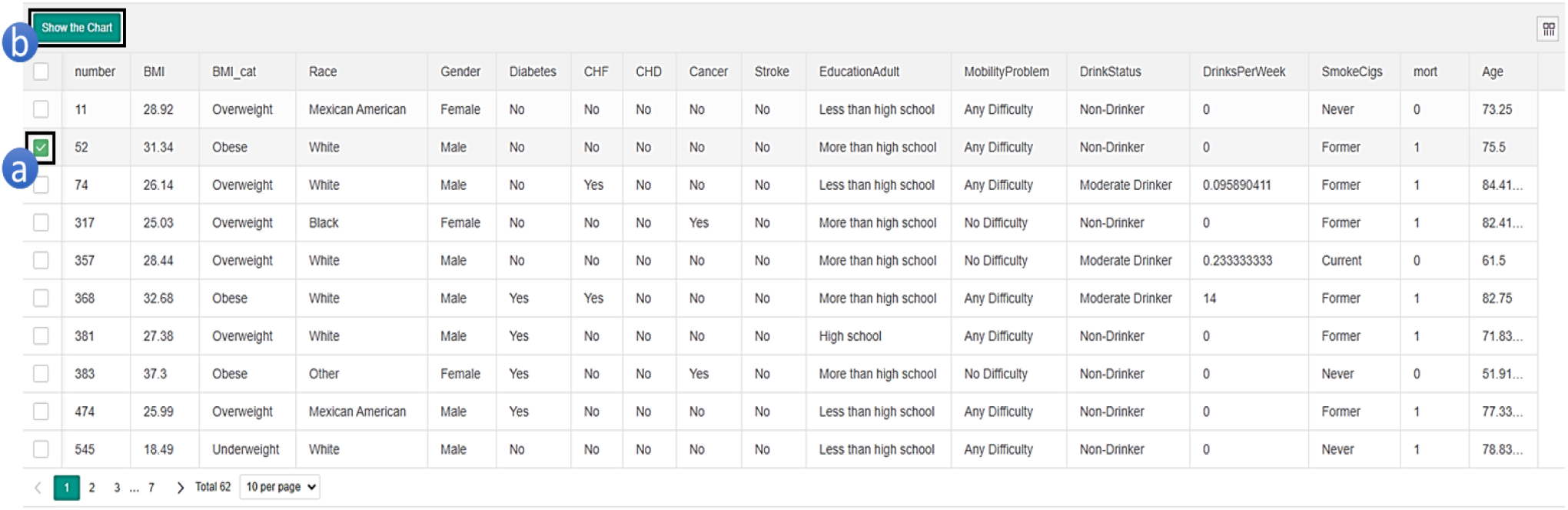
Sample matching results and corresponding clinical data

### 3.3 Explore Associations Between Hidden Layer Patterns and Observed Data

Our tool can facilitate the exploration between hidden layer patterns and observed data. Users can specify which co-variate or response variable to explore. Here we give two examples.

First, we can explore if there is any different pattern in the clinical variates among the selected samples with matched hidden layer patterns with the entire cohort. For instance, given the search criteria set <threshold = −0.15, time interval = (12, 13)>, we can display the gender ratio in both the matched results and the original sample set (Figure 5.a). As shown in Figure 5.a, in the original dataset, there are 1,454 males and 1,523 females, with a gender ratio of 1.04:1. However, there are 48 males and 29 females in our matched result, with a gender ratio of 1.65:1.

**Figure 5.**
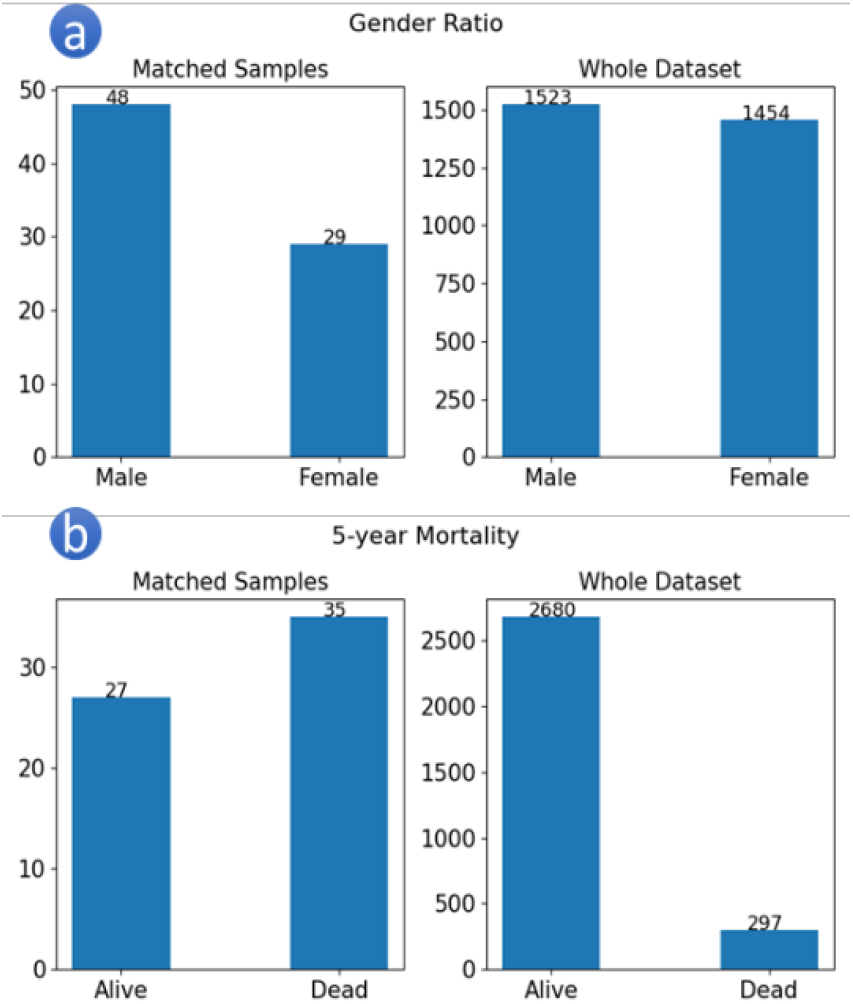
Two example exploration results

Another example is given the search criteria set <threshold =0.2, time interval = (12,12)>, display the mortality status in both the matched results and the original sample set (Figure 4 b). As shown in Figure 5.b, we can notice a vast difference where the mortality rate is 52% in the matched samples and 10% in the original dataset.

### 3.4 Explore the Relationship Between Covariates and Response Variables across Different Time Intervals

Beyond discovering the patterns among clinical covariates, we can further explore and visualize the relationship between time interval of interests and the response variable ---- 5-years mortality in this study. Here, we will make a hypothesis that the physical activity level early in the morning and late in the afternoon might be correlated to the participant’s mortality. So, we picked 4 intervals of interests listed in the table:

**Table 1.** Time intervals of interests

| No. | Time interval | Duration |
| --- | --- | --- |
| 1 | Tue 16:20 to Tue 19:40 | 3 hours 20 minutes |
| 2 | Thu 5:00 to Thu 11:40 | 6 hours 40 minutes |
| 3 | Sat 20:20 to Sat 23:40 | 3 hours 20 minutes |
| 4 | Sun 16:20 to Sun 19:40 | 3 hours 20 minutes |

The following explorations are based on the intervals listed above.

#### 3.4.1 Average Activity Count T-test

Now we want to know if the average activity counts in the time intervals above have a significant difference between living and deceased participants. First, our toolkit can plot the raw activity counts per living and deceased participants separately.

As shown in Figure 6, the line chart is the activity count for each participant during our first time interval of interest (Tue 16:20 to Tue 19:40), and each line represents the minute-level activity count for one individual. The chart with blue line on the top is the result for those participants who were alive in the 5 years following the study period, and the red stands for those deceased.

**Figure 6.**
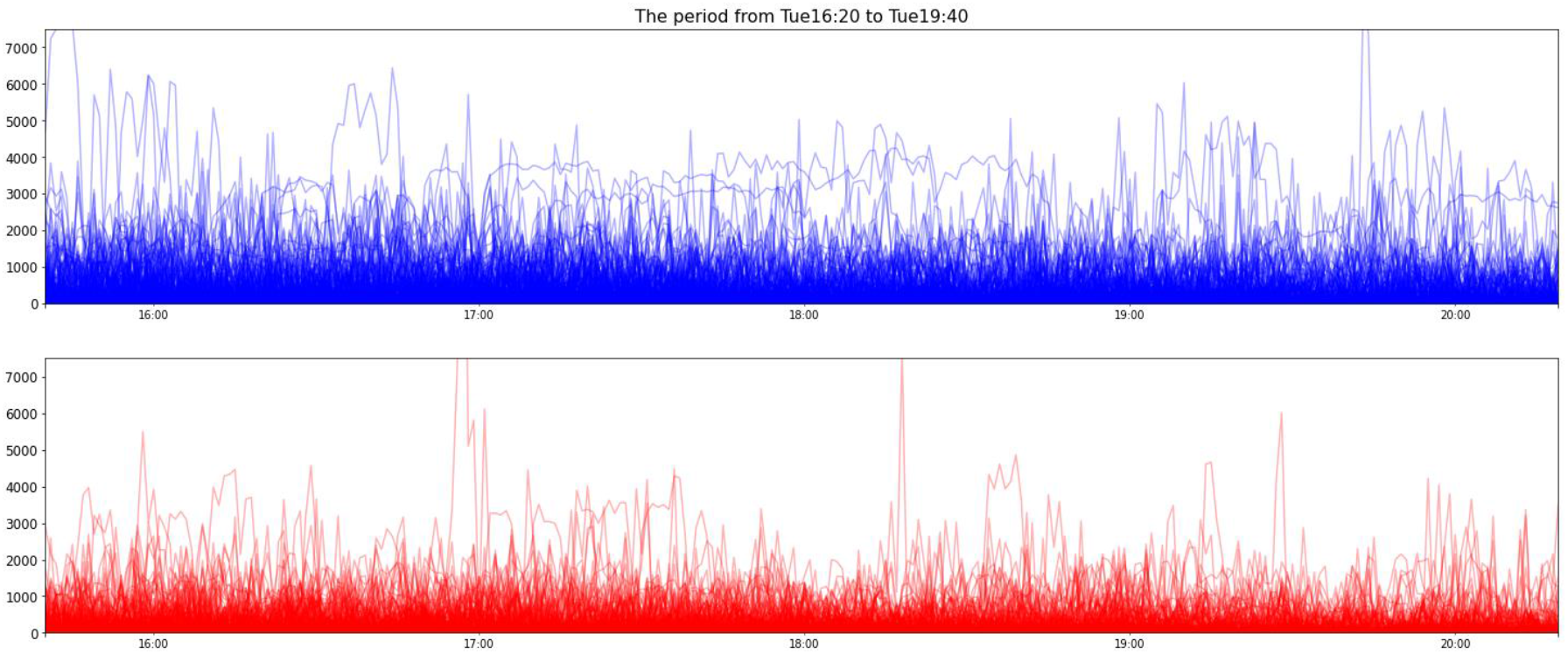
Activity Count from Tue 16:20 to Tue 19:40

Then, we operate a two-sample t-test to examine if the mean value of the two charts have significant differences. Our null hypothesis is the average activity counts per participant are equal between the alive and deceased. Since it turns out that all 4 time intervals’ average activity counts do not have homogeneity of variance, we conduct Welch’s test.

The Welch’s t-test result is recorded in Table 2.

**Table 2.** T-test result

| Time interval | Tue 16:20<br>to<br>Tue 19:40 | Thu 5:00<br>to<br>Thu 11:40 | Sat 20:20<br>to<br>Sat 23:40 | Sun 16:20<br>to<br>Sun 19:40 |
| --- | --- | --- | --- | --- |
| Mean count for alive | 220.46 | 202.00 | 83.96 | 196.24 |
| Mean count for dead | 128.76 | 130.14 | 54.23 | 120.43 |
| Welch's test p-value | 2.96e-6 | 1.61e-6 | 8.19e-4 | 1.16e-7 |

Set the significant value α = 0.05 and since all the p-values are less than our significant value, we can reject the null hypothesis. We can conclude there is significant difference in the average activity counts during these time intervals of interest between participants who were living or deceased in the five years following the study period.

#### 3.4.2 Sankey Plot

We are also interested in the activity transition pattern between participants who were alive or deceased after 5 years. Our logistic here is that given an activity threshold in each time interval of interest, we separate both those who were alive and deceased into active and nonactive groups. More specifically, we will have 4 following groups: alive-active, alive-nonactive, dead-active, dead-nonactive in each time interval of interest. The activity threshold is a hyperparameter that divides those participants into active and nonactive, and empirically, we pick the mean among all participants in the specific time interval as the threshold.

The Sankey Diagram is a type of flowchart that can easily present the transition between states in the time order. Our toolkit is able to present an interactive Sankey Diagram with PyEcharts package to show the transition pattern between active and nonactive among those alive or deceased separately.

As mentioned, those participants with mean activity counts larger than the average counts among all people in the same interval are considered active, and vice versa. Every column in Figure 7 represents the participants in one of our time intervals of interest, and each column of participants are grouped with their 5 years mortality status and active status during that time interval.

**Figure 7.**
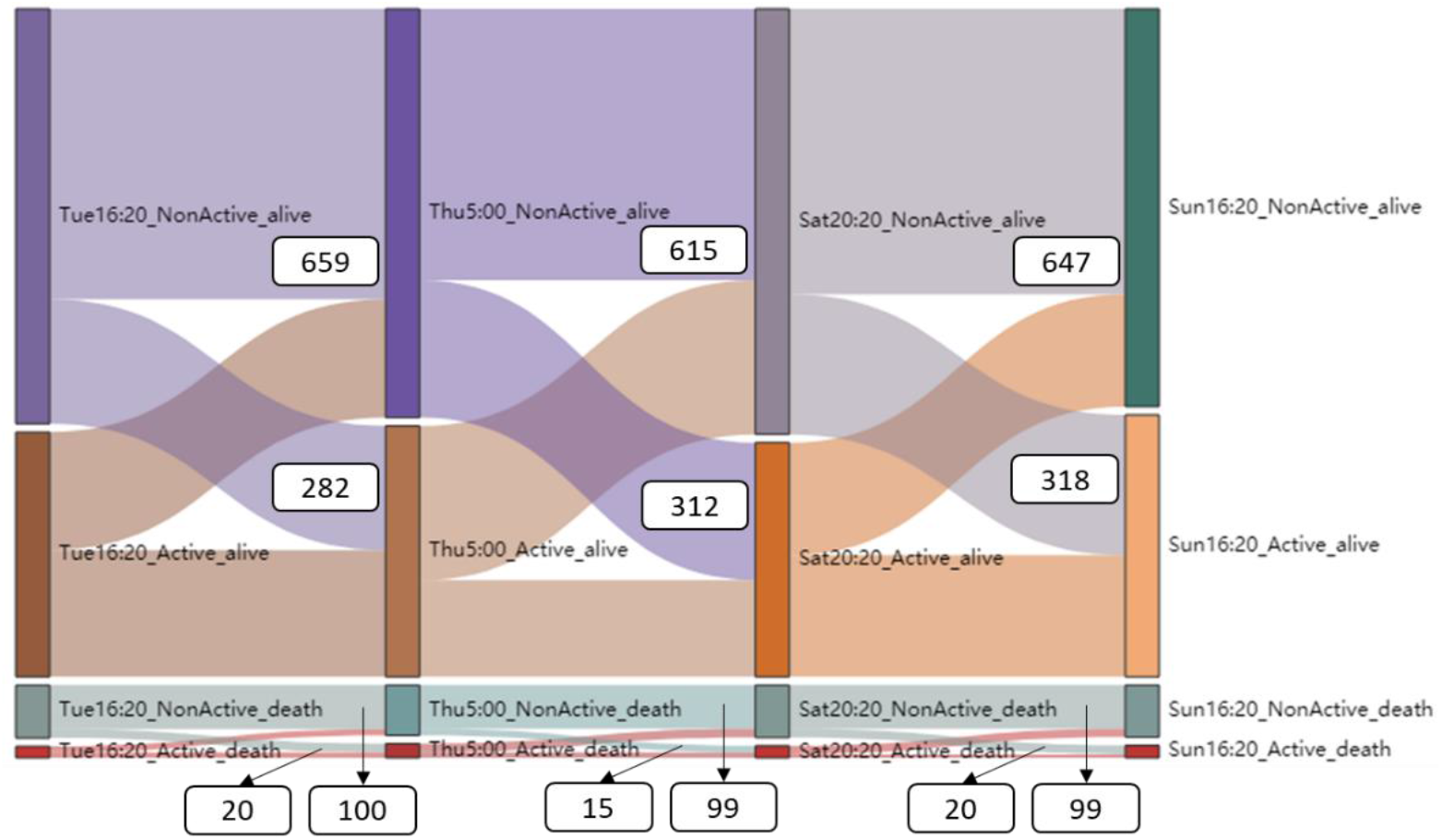
Sankey Diagram

In Figure 7, we annotated the population of alive and deceased respectively that transfer from nonactive state to active state between every two time intervals of interest. The values are collected below:

In Table 3, we calculated the transition rate from nonactive to active respectively in those alive and dead, and found that those participants who were alive after the 5-year time period had an overall higher transition rate from nonactive to active status (~0.3) compared to those who were deceased (~0.15). This pattern may be helpful to the 5 years mortality prediction model.

**Table 3.**
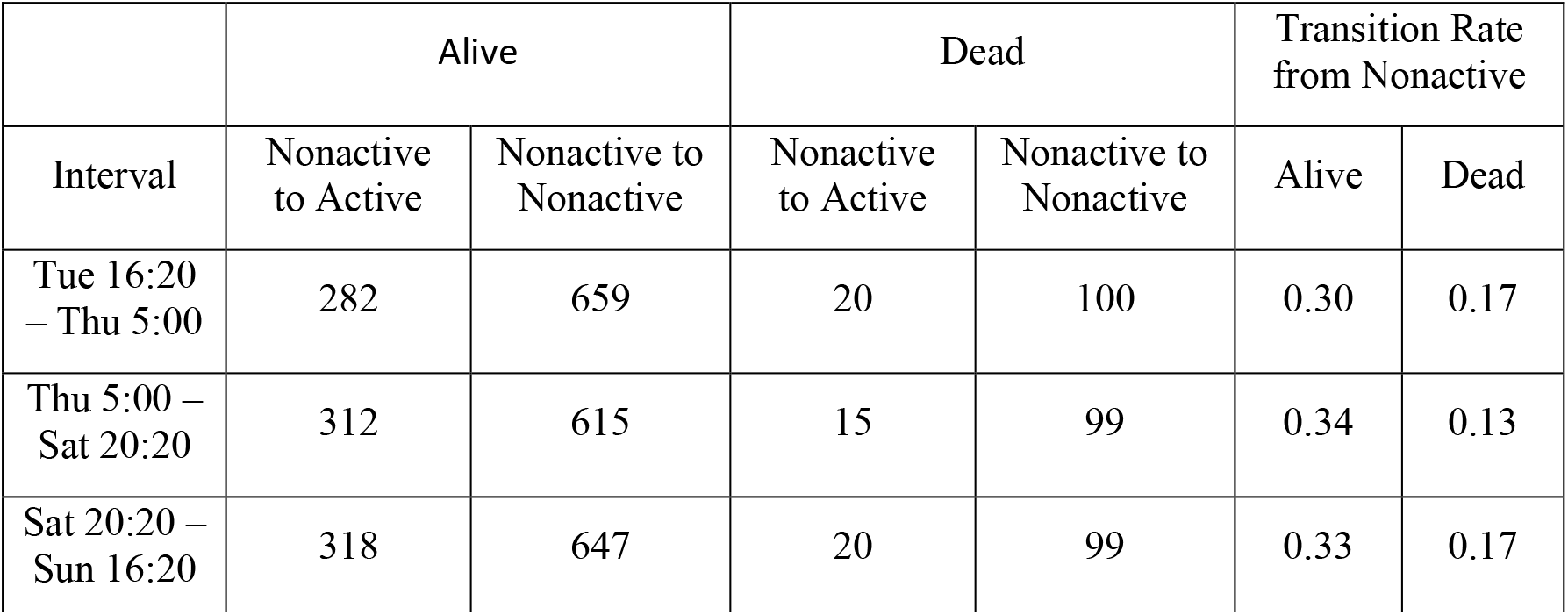
Alive and dead sample counts in 4 time intervals

## 4. Access to HiddenVis

HiddenVis is open source and available at https://github.com/jyan97/HiddenVis. HiddenVis is implemented based on Python, Flask and ECharts, and the environment requirement is listed in requirement.txt.

## 5. Conclusions and Discussions

In this paper, we developed a visualization web tool to facilitate the interpretation and exploration of RNN-based deep learning models, including LSTM, for time series data analysis. It can help researchers to see the values of hidden layers and fine-tune the visualization results: researches can highlight the hidden layer parameters by customized threshold and interval. They can also match the selected cohorts with similar patterns, explore their attributes, and can explore the relationship between two inputs.

A further direction to demonstrate the utility of HiddenViz is to benchmark it on larger cohorts and other DL models. However, both attempts require costly computational resources. In this project, we used the Google CoLab[11], and training one LSTM model with a fixed set of hyperparameters can take 40 minutes to 2 hours. Thus, fully fine-tuning a more complex model on larger cohorts may be cost-prohibitive at the moment. We do envision that such analysis will be feasible when GPU computation is more affordable.

The HiddenViz model is suitable for a wide range of DL-based accelerometer data analyses. It can be easily extended to the visualization and analysis of other temporal data. For example, an electrocardiogram (EKG) periodically reports electric signals[12]; the pulse oximeter monitor reports the oxygen saturation (SpO2) every second[13]. Researchers have been analyzing those data using DL models. Thus HiddenViz can be also applied to interpret the DL models, reveal important clinical cohorts, and associate versatile types of temporal data to health outcomes.

## References

[1] Qin, Yao, et al. “A dual-stage attention-based recurrent neural network for time series prediction.” Proceedings of the 26th International Joint Conference on Artificial Intelligence. 2017.

[2] Selvin, Sreelekshmy, et al. “Stock price prediction using LSTM, RNN and CNN-sliding window model.” 2017 international conference on advances in computing, communications and informatics (icacci). IEEE, 2017.

[3] Choi, Edward, et al. “Doctor ai: Predicting clinical events via recurrent neural networks.” Machine Learning for Healthcare Conference. 2016.

[4] Sun, Fei, et al. “BERT4Rec: Sequential recommendation with bidirectional encoder representations from transformer.” Proceedings of the 28th ACM International Conference on Information and Knowledge Management. 2019.

[5] Lyons, G. M., et al. “A description of an accelerometer-based mobility monitoring technique.” Medical engineering & physics 27.6 (2005): 497–504.

[6] Troiano, Richard P., et al. “Physical activity in the United States measured by accelerometer.” Medicine and science in sports and exercise 40.1 (2008): 181.

[7] Lohne-Seiler, Hilde, et al. “Accelerometer-determined physical activity and self-reported health in a population of older adults (65–85 years): a cross-sectional study.” BMC Public Health 14.1 (2014): 284.

[8] Smirnova, Ekaterina, et al. “The predictive performance of objective measures of physical activity derived from accelerometry data for 5-year all-cause mortality in older adults: National Health and Nutritional Examination Survey 2003–2006.” The Journals of Gerontology: Series A 75.9 (2020): 1779–1785.

[9] Chalapathy, Raghavendra, Ehsan Zare Borzeshi, and Massimo Piccardi. “Bidirectional LSTM-CRF for clinical concept extraction.” arXiv preprint arXiv:1611.08373 (2016).

[10] Bang, Seojin, et al. “Explaining a black-box using deep variational information bottleneck approach.” arXiv preprint arXiv:1902.06918 (2019).

[11] Bisong, Ekaba. “Google Colaboratory.” Building Machine Learning and Deep Learning Models on Google Cloud Platform. Apress, Berkeley, CA, 2019. 59–64.

[12] Al Rahhal, Mohamad Mahmoud, et al. “Deep learning approach for active classification of electrocardiogram signals.” Information Sciences 345 (2016): 340–354.[9]

[13] Jensen, Louise A., Judee E. Onyskiw, and N. G. N. Prasad. “Meta-analysis of arterial oxygen saturation monitoring by pulse oximetry in adults.” Heart & lung 27.6 (1998): 387–408.

